# Context-aware single-cell multiome approach identified cell-type specific lung cancer susceptibility genes

**DOI:** 10.1101/2023.09.25.559336

**Authors:** Erping Long, Jinhu Yin, Ju Hye Shin, Yuyan Li, Alexander Kane, Harsh Patel, Thong Luong, Jun Xia, Younghun Han, Jinyoung Byun, Tongwu Zhang, Wei Zhao, Maria Teresa Landi, Nathaniel Rothman, Qing Lan, Yoon Soo Chang, Fulong Yu, Christopher Amos, Jianxin Shi, Jin Gu Lee, Eun Young Kim, Jiyeon Choi

## Abstract

Genome-wide association studies (GWAS) identified over fifty loci associated with lung cancer risk. However, the genetic mechanisms and target genes underlying these loci are largely unknown, as most risk-associated-variants might regulate gene expression in a context-specific manner. Here, we generated a barcode-shared transcriptome and chromatin accessibility map of 117,911 human lung cells from age/sex-matched ever- and never-smokers to profile context-specific gene regulation. Accessible chromatin peak detection identified cell-type-specific candidate *cis*-regulatory elements (cCREs) from each lung cell type. Colocalization of lung cancer candidate causal variants (CCVs) with these cCREs prioritized the variants for 68% of the GWAS loci, a subset of which was also supported by transcription factor abundance and footprinting. cCRE colocalization and single-cell based trait relevance score nominated epithelial and immune cells as the main cell groups contributing to lung cancer susceptibility. Notably, cCREs of rare proliferating epithelial cell types, such as AT2-proliferating (0.13%) and basal cells (1.8%), overlapped with CCVs, including those in *TERT*. A multi-level cCRE-gene linking system identified candidate susceptibility genes from 57% of lung cancer loci, including those not detected in tissue- or cell-line-based approaches. cCRE-gene linkage uncovered that adjacent genes expressed in different cell types are correlated with distinct subsets of coinherited CCVs, including *JAML* and *MPZL3* at the 11q23.3 locus. Our data revealed the cell types and contexts where the lung cancer susceptibility genes are functional.

## Introduction

Lung cancer is the leading cause of cancer deaths and affects diverse populations worldwide^1^. Although the major risk factor for lung cancer is tobacco smoking, up to 25% of lung cancers arise in never-smokers^2^. Lung cancer heritability is estimated as ∼8-20%^3–6^, and genome-wide association studies (GWAS) have identified over 50 risk loci so far. Notably, a substantial proportion of these loci are specifically observed in subgroups based on histological subtypes, ancestry, and smoking status^7–9^, suggesting heterogeneous genetic mechanisms of lung cancer susceptibility. While several biological pathways (e.g., telomere biology, immune response, DNA damage repair) have been highlighted from these GWAS loci by single-locus-based studies^8^, functional variants and susceptibility genes are uncharacterized from most lung cancer GWAS loci.

Identifying target genes from a GWAS locus is challenging, because most of the risk-associated variants are in non-protein-coding regions and do not alter amino acid sequences^10,11^. Instead, most GWAS variants appear to function as *cis*-regulatory elements (CREs) and alter gene expression levels by affecting the binding of *trans*- acting factors (e.g., transcription factor, TF) in gene promoters and enhancers^12^. Consistent with this hypothesis, approaches linking genetic variants to their putative targets of transcriptional regulation have been powerful in identifying susceptibility genes from GWAS loci. For example, expression quantitative trait loci (eQTL) approach can detect the association between GWAS variants and transcript levels of nearby genes. Chromatin interaction approach can link GWAS variants located in candidate CREs (cCREs) to target gene promoters, using physical interaction via chromatin looping^13^. However, the regulatory activities of the CREs are highly specific to cell types and cellular contexts^14^, and the current bulk tissue-based eQTL and cell-line-based chromatin interactions data might not comprehensively capture these diverse biological contexts as well as the effect of environmental exposures on gene expression regulation.

Lung tissue has diverse cell types, and lung tumorigenesis is considered an interplay between the cells of cancer origin and their microenvironment, including immune cells, in the presence of external exposures such as smoking. Cells of lung cancer origin have been mainly traced in animal models, which suggested the roles of alveolar type II (AT2) and club cells for lung adenocarcinoma, basal cells for lung squamous cell carcinoma, and neuroendocrine cells for small cell lung cancer, which are all in the epithelial group of lung cells. Recent single-cell approaches using human and mouse lung tissues identified transient states of AT2 cells considered as stem/progenitor cells^15–19^, hinting at their potential involvement in lung adenocarcinoma. However, many of these epithelial cell types of presumed cancer origin are rare in lung tissue (<5%) based on multiple classical studies^15^. Furthermore, in fresh-dissociated single-cell suspension of lung tissue, epithelial cells tend to be depleted while immune cells are overrepresented^15^. Given this complexity, even when bulk-tissue-based eQTL identified multiple candidate target genes, it is difficult to determine in which cell types (e.g., cells of cancer origin vs. immune cells) the lung cancer susceptibility genes might be functional. Moreover, complex interplay of multiple target genes potentially regulated by one or more risk-associated variants in different contexts has not been explored.

To this end, single-cell multiome approaches combining single-nucleus RNA-sequencing (snRNA-seq) and single-nucleus assay for transposase-accessible chromatin with sequencing (snATAC-seq) can provide a powerful alternative to annotating GWAS loci^20,21^. While there are abundant single-cell based resources for human lung tissues as recently compiled by Human Lung Cell Atlas^18^, single-cell profiling of chromatin accessibility in human lung tissues is still limited to a small number of datasets^22^ and to our knowledge a joint profiling of transcriptome and chromatin accessibility in the same barcoded cells has not been performed. Moreover, cell-type specific gene expression changes by lung cancer-relevant exposures such as smoking have not been formally profiled in human lung tissues. In this study, we performed a barcode shared snRNA-seq and snATAC-seq of human lung tissues while incorporating age/sex-matched Korean ever- and never-smokers and adopting a strategy to enrich epithelial cell population. This multiome approach enables joint clustering between the two modalities, identification of cell-type specific cCREs, and linkage of cCREs to target genes^23^. By capturing the lung cancer-relevant cellular contexts, we extend the functional characterization of lung cancer GWAS loci representing diverse populations.

## Results

### Single-cell multiome design representing lung-cancer-relevant contexts

To generate a single-cell dataset representing lung-cancer-relevant cellular contexts, we designed a study focusing on smoking status as the main environmental exposure and cells of lung cancer origin to capture the endogenous lineage-specific gene regulation. We collected tumor-distant normal lung tissues from sex- and age-matched ever- and never-smokers (n = 8 each), which were fresh-dissociated and cryopreserved before further processing (**Figure 1A**; **Methods**). All the baseline characteristics of these samples are presented in **Table S1**. Our design adopted barcode-shared snATAC-seq and snRNA-seq to jointly profile chromatin accessibility and gene expression from each cell type of lung tissues (**Figure 1B**). This approach allows variant-cCRE colocalization, cCRE-gene linkage, and characterization of lung cancer GWAS loci at the levels of variant, gene, and cell type (**Figure 1C-D**). Given that immune cells tend to be overrepresented in normal lung tissues^15^ and epithelial cells are more vulnerable to dissociation process as well as freezing and thawing, we employed balancing of the lung cell groups using cell surface marker labeling followed by fluorescence-activated cell sorting (FACS) (**Methods**). Antibodies against EPCAM, CD31, and CD45 were used to sort the live single-cells into four major categories: “epithelial” (EPCAM^+^CD45^−^CD31^−^), “immune” (EPCAM^−^CD45^+^CD31^−^), “endothelial” (EPCAM^−^CD45^−^CD31^+^), and “stromal” (EPCAM^−^CD45^−^CD31^−^) (**Figure 1E**). To enrich epithelial cells, which include cell types considered relevant for lung cancer origin, we collected maximum number of “epithelial” cells from each sample while keeping a substantial ratio of “immune”, “endothelial”, and “stromal” groups (6:3:1 mixing of presumed “epithelial”, “immune”, and the remaining fractions) (**Figure 1E; Methods**). Based on flow cytometry, our strategy resulted in 20.3-52.4% (median 38.8%) of estimated “epithelial” cells across 16 samples after enrichment, which is a significant improvement (∼4-fold increase) compared to 4.3-23.6% (median 10.0%) before balancing (**Figure 1F**). “Immune” (median 31.4%), “endothelial” (median 25.3%), and “stromal” cells (median 4.2%) were also estimated at expected proportions after balancing.

**Figure 1.**
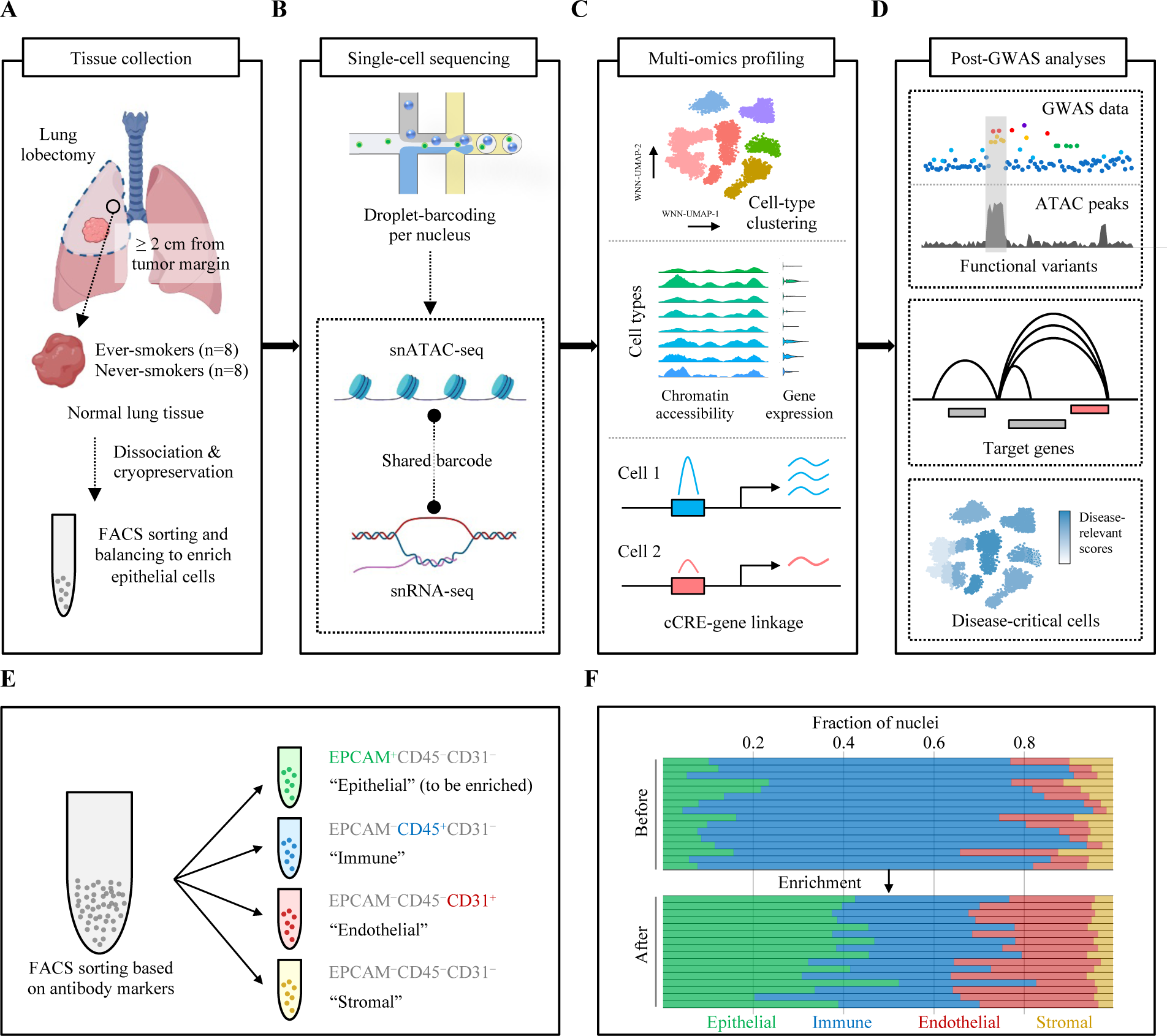
Graphic summary of study design, workflow, and enrichment strategy. Overview of the study pipeline, including tissue collection **(A)**, single-cell sequencing **(B)**, multiome profiling/analyses of chromatin accessibility/gene expression **(C)**, and post-GWAS analyses of functional variants, target genes, and disease-relevant cells underlying lung cancer susceptibility loci **(D)**. **(E)** Antibody markers of EPCAM, CD31, and CD45 were used to sort the live single-cells: EPCAM^+^CD45^−^CD31^−^ (“epithelial”), EPCAM^−^CD45^+^CD31^−^ (“immune”), EPCAM^−^CD45^−^CD31^+^ (“endothelial”), and EPCAM^−^CD45^−^CD31^−^ (“stromal”). Epithelial cells were enriched by balancing the ratios among “epithelial”, “immune”, “endothelial”, and “stromal”. **(F)** Bar plot shows the nuclei fraction of “epithelial” (green), “immune” (blue), “endothelial” (red), and “stromal” (yellow) before and after enrichment. Each bar represents an individual sample following the order of sample IDs listed in **Table S1**.

### Simultaneous profiling of single-nucleus chromatin accessibility and gene expression identified classic and transient lung cell types

Using the balanced lung cells, we performed nuclei isolation followed by barcode-shared snATAC-seq and snRNA-seq to generate single-nucleus chromatin accessibility and gene expression matrices (**Methods**). The detailed summary metrics of the sequencing results are provided in **Table S2**. After filtering low-quality nuclei, likely empty droplets, and doublets (**Methods**), we obtained 117,911 single-nuclei from 16 samples with high-quality chromatin accessibility and gene expression profiles (**Table S3**). After correcting the batch effects, these nuclei were clustered and assigned into 23 cell types (**Figure 2A, Table S4**) based on both modalities (**Methods**). They represent epithelial (AT2, alveolar type I or AT1, club, ciliated, goblet, basal, AT1/AT2, and AT2-proliferating), immune (natural killer or NK, T, macrophage, monocytes, B, dendritic, and NK/T), endothelial (lymphatic, artery, vein, and capillary), and stromal (fibroblasts, smooth muscle, mesothelial, and myofibroblasts) cells. A total of 47,453 epithelial cells (40.2%) were identified (**Figure 2B**), which closely matched the estimation based on flow cytometry, supporting the validity of our enrichment strategy and cell type annotation. We identified 36,308 immune (30.8%), 28,395 endothelial (24.1%), and 5,755 stromal cells (4.9%) (**Figure 2B**), and all 23 cell types were detected in 14 or more samples (**Table S5**), supporting that the cell-type identity is not driven by a single or a few samples with batch effects. Consistent with published studies of human lung, we observed cell-type-specific canonical gene expression markers for all 23 cell types (**Figure 2C**, additional markers in **Figure S1**). Using snATAC-seq data of these 23 cell types, we detected a total of 330,453 chromatin accessibility peaks (cCREs). Similar to the observations in previous snATAC-seq and multiome studies in different tissue types^20,24,25^, a substantial proportion (37%) of these cCREs were detected only in a single cell type (**Figure S2A**). Most of these cCREs were located in genic or gene promoter regions, including promoter (22%), exonic (4.3%), and intronic regions (46.1%) (**Figure S2B**). The cCREs detected in all 23 cell types displayed a higher proportion of promoter-located cCREs (91.8%) compared to those in the cell-type private ones (15.9%). Conversely, cell-type private cCREs displayed a higher proportion of intronic (48.4%) and intergenic cCREs (29.6%), which is consistent with the roles of distal enhancers in cell-type specific gene regulation^24^. Notably, we identified a rare cell type, AT2-proliferting cells (0.13%), which shows markers of both AT2 (*SFTPD*) and club cells (*SCGB3A2*) but specifically expresses the markers of cell proliferation, *STMN1*, *TYMS*, *TOP2A*, *CDK1*, and *MKI67* similar to two previous studies (“AT2-proliferating” and “cycling-AT2”)^18,19^ (**Figure 2D**). Consistent with the gene expression data, we observed distinct chromatin accessibility around the promotor-adjacent enhancer regions of *STMN1*, *TYMS*, *CDK1*, and *MKI67* in AT2-proliferating cells when compared to those in AT2 or club cells. These data highlighted unique values of epigenomic data supplementing transcriptome data in navigating cell-type specific gene regulation (**Figure 2D**).

**Figure 2.**
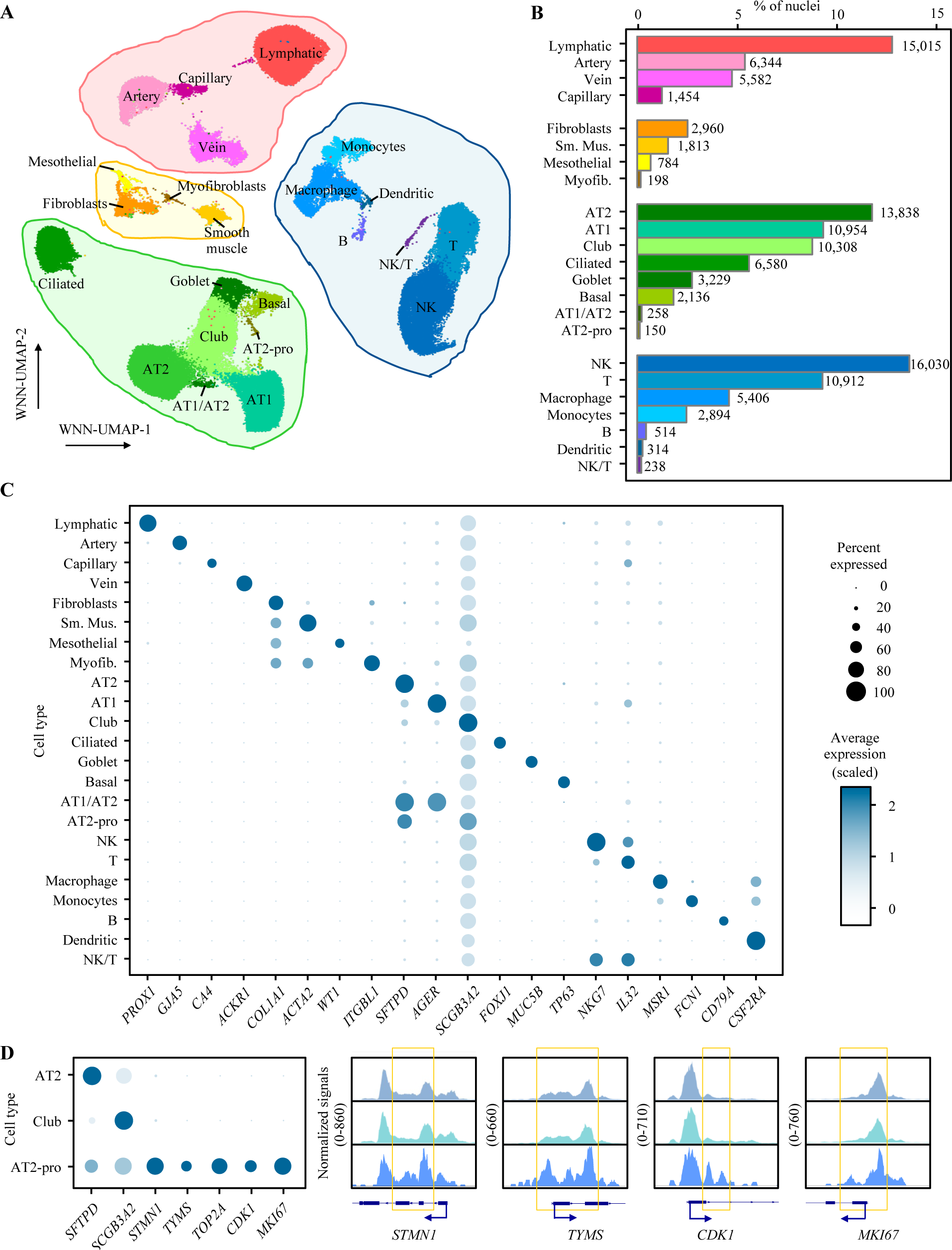
Identification of major cell types of the human lung via joint profile of snRNA-seq and snATAC-seq. **(A)** Weighted Nearest Neighbor (WNN) clustering results of the 117,911 single-nucleus using the profiles of chromatin accessibility and gene expression after quality controls with cell-type annotation based on canonical markers, which resulted in 23 cell types across endothelial (red shades), stromal (yellow shades), epithelial (green shades), and immune (blue shades) cells. **(B)** Fraction of nuclei of each cell type relative to the total number of nuclei. Numbers next to each bar denote absolute counts out of 117,911 nuclei. **(C)** Dot plot visualizing the normalized RNA expression of selected marker genes by cell type. The color and size of each dot correspond to the scaled average expression level and fraction of expressing cells, respectively. Additional markers are shown in **Figure S1**. **(D)** The dot blot on the left visualizes the normalized RNA expression of AT2, club, and AT2-proliferating cells (AT2-pro) with the same style as panel C. The sequencing tracks on the left visualize chromatin accessibility signals around selected marker genes by cell type. Each track represents the aggregate snATAC signal of all three included cell types normalized by the total number of reads in the transcription start site (TSS) region. Arrows show the transcriptional directions of the genes. Coordinates for each region are as follows: *STMN1* (chr1:25904993-25908229), *TYMS* (chr18:657083-658911), *CDK1* (chr10:60768638-60777926), *MKI67* (chr10:126261833-126268756). Yellow squares highlight the main differences of chromatin accessibility among three cell types.

### Smoking affects intercellular communications of human lung

Among 117,911 total nuclei, 59,250 were from ever-smokers and 58,661 were from never-smokers. We first compared the gene expression between ever-and never-smokers in each cell type to nominate the “smoking-responsive genes” using a stringent pseudobulk method (**Methods**). While only one gene (*AC017002.5*, FDR = 0.0066, lymphatic cells) passed the multiple testing correction cutoff (FDR < 0.05), 24 genes displayed suggestive differences at a relaxed cutoff (P < 1×10^-4^), including previously implicated genes in smoking and related traits (**Table S6**).

We then asked whether cell-cell communication is different based on smoking status, since smoking has been associated with cellular processes that involve intercellular interactions such as inflammation and immune functions^34^. Based on the known ligand-receptor pairs in CellChatDB, cell-cell communication strengths were summarized at the pathway level and compared between the cells from ever- and never-smokers (**Methods**). We nominated the top five pathways displaying the largest differences in the communication strength based on smoking status across all cell types. The top five pathways with elevated communication strength in ever-smokers (MHC-I, UGRP1, CD46, NRXN, NEGR) and in never-smokers (MHC-II, VCAM, CD6, ALCAM, THBS) displayed the differences both at the level of individual cell types and across all cell types (**Figure S3**). Notably, Major Histocompatibility Complex (MHC)-I and -II pathways exhibited an opposite trend across all cell types between ever- and never-smokers. This observation is, in principle, consistent with a recent study reporting that the levels of *HLA-A, B, C* (MHC-I) were higher in the lung adenocarcinoma cells from smokers compared to never-smokers, and MHC-II genes were elevated in cell subclusters representing never-smoker cancer cells^33^. MHC-II genes (*HLA-DRA*, *HLA-DRB1*) were also suppressed by smoking in multiple types of airway epithelial cells in a previous study^29^. Our findings validated previous observations based on gene expression levels and extended the understanding to the cell-cell communication across the cell types of lung.

### Epithelial and immune cell-specific *cis*-regulation underlies lung-cancer-associated variants

Using the cCREs that we detected in each cell type of lung, we aimed to better understand the risk loci identified by GWASs of lung cancer. Combining the four most recent lung cancer GWASs representing European, East Asian, and African ancestries as well as major lung cancer histological types and smoking status^9,35–37^, we complied a set of candidate causal variants (CCVs) (**Figure 3A**). We included 51 non-overlapping GWAS loci (**Table S7**) and 2,574 unique CCVs (**Table S8**) based on the GWAS statistics and LD (log likelihood ratio 1:1000 or *R*^2^ ≥ 0.8 to the lead variant in the study population). To characterize how these CCVs align with cCREs, we performed CCV-cCRE colocalization followed by allelic TF abundance and footprinting analyses (**Figure 3B**).

**Figure 3.**
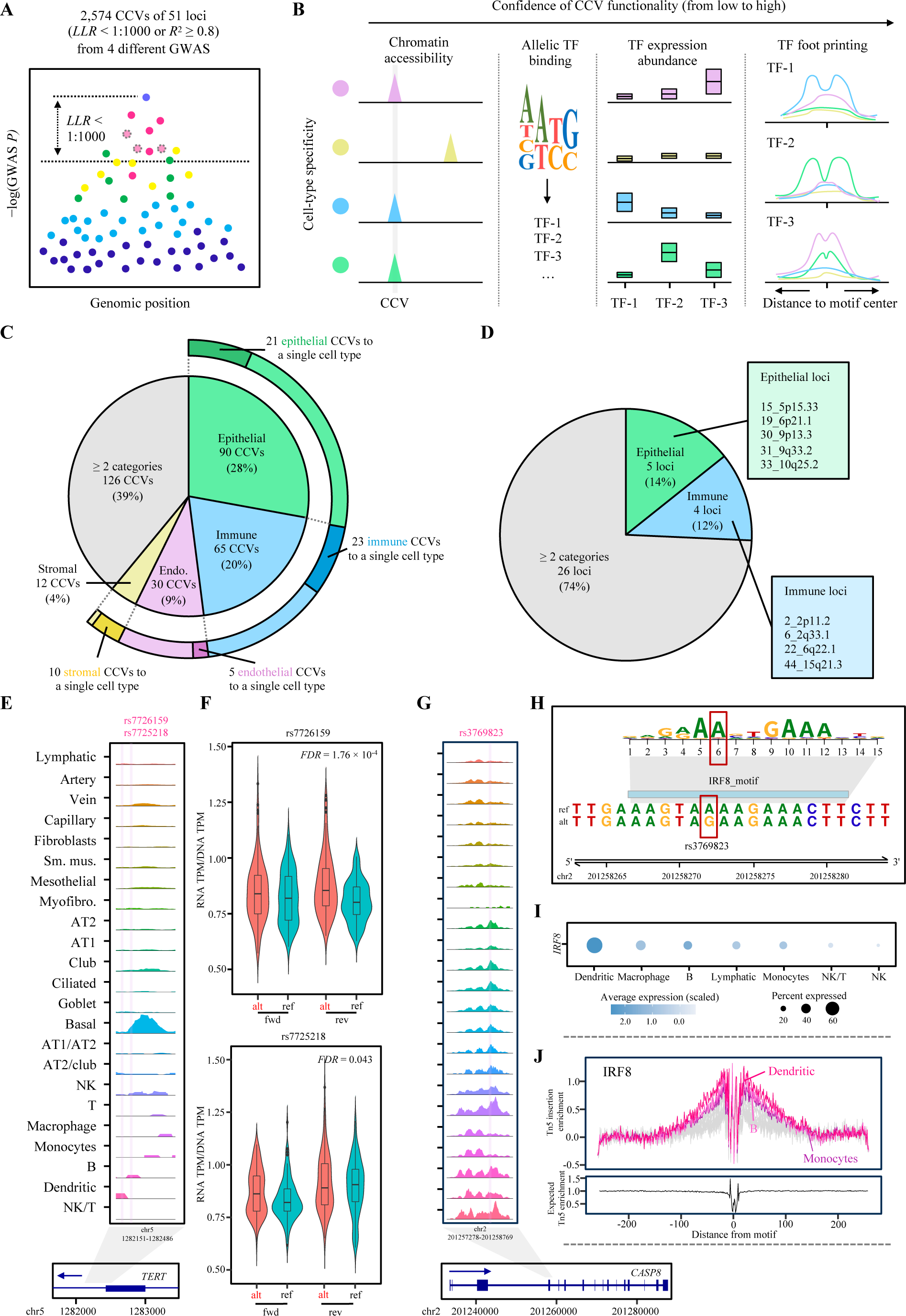
Characterization of the cell-type specificity underlying lung-cancer-associated functional variants. **(A)** Illustration of the candidate causal variants (CCVs) selection from GWAS loci. Each dot represents a variant, and the linkage disequilibrium (LD) to lead variant is color-coded. Dots with dashed outline represent the variant in high-LD with the lead variant but not covered in the original GWAS summary statistics. **(B)** The confidence level of the CCV functionality is established by colocalizing them with accessible chromatin regions, predicting allelic TF binding effects, assessing TF abundance, and TF footprinting. **(C)** Piechart presents the fraction of CCVs assigned to different cell-type categories (inner pie) and assigned to a single cell type (external pie). **(D)** Piechart presents the fraction of loci assigned to different categories. **(E)** The sequencing tracks representing chromatin accessibility are displayed. The rsIDs above the tracks are marked with vertical pink lines to indicate their positions. Each track represents the aggregated snATAC signal of all cell types, normalized by the total number of reads in the regions (y-axis scale 0-380). Arrow depicts the transcriptional direction of *TERT*. (**F**) The transcriptional activity of 145 bp sequences encompassing rs7726159 (upper) and rs7725218 (lower) were tested in A549 lung cells. The activity is presented as the RNA TPM/DNA TPM from massively parallel reporter assays (MPRA). Both alleles (alternative, alt or lung cancer risk-associated; reference, ref) are shown in forward (fwd) and reverse (rev) directions. Center lines show the medians; box limits indicate the 25th and 75th percentiles; whiskers extend 1.5 times the interquartile range from the 25th and 75th percentiles, outliers are represented by dots. Density is reflected in the width of the shape. FDR values were calculated by the Wald test and corrected by the Benjamini-Hochberg procedure. (**G**) The sequencing tracks of chromatin accessibility, CCVs, and gene transcriptional directions using the same style as the left part of panel E (y-axis scale 0-490). (**H**) The upper part displays the position weight matrix (PWM) as the height of the motif logos. The position of the variant (rs3769823) within the motif is indicated with a red box. The lower part shows the genomic region containing the IRF8 motif and allelic binding site of rs3769823. (**I**) The normalized mRNA expression of *IRF8* is displayed as the dot plots across 7 cell types with the highest *IRF8* expression. The color and size of each dot corresponds to the scaled average expression level and fraction of cells expressing *IRF8*, respectively. (**J**) The upper part displays the footprint analysis of the IRF8 across cell types. The top three cell types (color-coded) showing IRF8 footprint are dendritic, B, and monocytes. The remaining cell types are in gray. Footprints were corrected for Tn5 insertion bias by subtracting the Tn5 insertion signal from the footprinting signal. The lower part shows the expected Tn5 enrichment based on distance from motif.

First, we observed that 323 CCVs (12.5% of the tested CCVs) of 35 GWAS loci (68.6% of the GWAS loci) colocalized with a cCRE from one or more cell types (**Table S9**), which indicated that a substantial proportion of GWAS loci was covered by this approach and a considerable variant prioritization was achieved. To nominate the lung cell types where lung cancer GWAS signals are enriched the most, we first integrated GWAS and snATAC-seq data using SCAVENGE^38^. SCAVENGE directly uses GWAS variants that colocalize with cCREs as opposed to indirectly linking the variants to gene expression data based on the distance to nearby genes or bulk-based expression correlation. We calculated the trait relevance scores (TRS) for each cell and compared them between cell types and cell categories (**Methods**). At the level of cell categories, epithelial and immune cell types showed higher mean TRS compared to endothelial and stromal cell types (3.60 and 3.57 vs 2.71 and 2.84, respectively) (**Figure S4**). Notably, AT2 cells and AT2-proliferating cells displayed the highest mean TRS, which is consistent with a previous study pointing to AT2 cells for showing a suggestive enrichment of lung adenocarcinoma susceptibility based on lung scRNAseq data^18^. We then defined the cell-type specificity for each CCV-colocalized cCRE by assessing whether the cCRE were being exclusively called in specific cell types or categories (**Methods**). We noted that 61% of the CCV-colocalized cCREs were assigned to a single cell-type category, with 28% to epithelial, 20% to immune, 9% to endothelial, and 4% to stromal (**Figure 3C**). For example, in loci such as 30_9p13.3 and 19_6p21.1, we observed CCV-colocalizing cCREs that were mainly detected in epithelial category and in 6_2q33.1 and 5_2p23.3, those mainly observed in immune category (**Figure S5**). There were substantial proportions of CCVs that were assigned to a single cell type (**Figure 3C**). At locus level, we observed 5 loci that were assigned to epithelial category and 4 that were assigned to immune category (**Figure 3D**). Notably, 17 CCV-colocalized cCRE from 12 loci were detected in AT2-proliferating cells, suggesting the role of transient epithelial cell types in lung cancer susceptibility. These results suggested that epithelial and immune cells are important cell types underlying the lung cancer GWAS loci.

Among the cell-type specific CCV-colocalizing cCREs were those from rare lung epithelial cell types. In the locus 15_5p15.33, two CCVs (rs7726159 and rs7725218) colocalized with a cCRE within *TERT* which is specific to basal cells, an epithelial cell type accounting for 1.8% of the total lung cells in our datset (**Figure 3E**). *TERT* expression is active in embryonic stem cells but silenced in differentiated cells; re-activation of *TERT* is central to cellular immortalization and tumorigenesis in multiple cancer types^39^. Consistent with this idea, basal cells in the epithelium of airway and lung are considered stem/progenitor cells with self-renewal ability^40^. To further examine the function of these two cCRE-overlapping CCVs, we performed reporter assays comparing the enhancer activity of lung-cancer risk and protective alleles of each variant within a short sequence (145bp) using A549 lung cancer cell line (**Methods**). The results demonstrated that one variant displayed significant allelic transcriptional activity at false discovery rate (FDR) < 1% with the lung cancer risk-associated allele showing higher levels, and the other showed a similar trend (FDR = 0.043) (**Figure 3F**). These data suggested that these lung cancer CCVs might exert their function through the basal cell specific cCRE of lung, highlighting the utility of our approach in elucidating cell-type specific gene regulation underlying lung cancer susceptibility.

To further prioritize potential functional variants from the cCRE-colocalized variants, we performed TF analyses using cell-type-specific TF expression and cCRE features. We first predicted TF binding affinity of each CCV (**Figure 3B**), and 50% of the variants displayed allelic differences in binding scores to one or more TFs (**Table S9**). For these allelic TFs, we investigated cell-type specific expression levels. Fifty-six allelic TFs were abundantly expressed in the same cell type(s) where the cCREs were detected (**Table S9**; **Methods**). We further performed TF footprinting for all the allelic TFs (**Methods**), and 82 TFs exhibited an enriched accessibility to their motif-flanking regions within cCREs. Overall, 111 CCVs from 29 GWAS loci displayed either a TF footprint or cell-type matching TF abundance for a predicted allelic-binding TF. Among them was a previously identified multi-cancer-associated functional variant, rs3769823, colocalized with a multi-cell-type cCRE within a *CASP8* alternative promoter (**Figure 3G**). This missense variant was shown to alter CASP8 protein activity and affect apoptosis and proliferation of lung cancer cells^41^ but also displayed a strong allelic transcriptional activity in immortalized melanocytes and melanoma cells^42^. Among many TFs predicted to display allelic binding affinity to this variant, IRF8 was abundantly expressed in dendritic cells, a cell type where the footprints of its binding motif were also detected (**Figure 3H-J**; **Table S9**). Identification of a known functional variant provided support for our TF abundance and footprinting approach for variant prioritization as well as further insights into potential cell-type specific roles of a known susceptibility gene. Our dataset identified lung cancer-relevant cell types of lung and nominated lung cancer CCVs that might be under cell-type specific regulation, including a subset involving potential allelic TF binding.

### Multi-level linkage identified context-specific candidate susceptibility genes from lung cancer loci

We leveraged the barcode-matched multiome data to identify target genes of the lung cancer CCV-colocalized cCREs by employing a multi-level cCRE-gene linkage (**Methods**). Specifically, we first performed a “cCRE module” analysis to find a group of cCREs displaying co-accessibility and assign them a unique cCRE module membership. We then performed cCRE-cCRE and cCRE-gene correlation analyses to identify the cCREs or cCRE modules whose accessibility is significantly correlated with the accessibility of a promoter cCRE and/or the expression of a gene in *cis* (+/- 1Mb) (**Methods**, **Figure 4A**). We reasoned that a direct correlation between a cCRE accessibility and a gene expression provides the strongest evidence compared to a link through a highly correlated cCRE (co-accessible score > 0.5) or a cCRE module membership (co-accessible score > 0.32). We also considered the location of the cCREs within a gene promoter (distance to a transcription start site, TSS) or when a cCRE or a cCRE module is correlated with a promoter cCRE without direct evidence of expression level correlation albeit at a lower priority level. This multi-level system provided six different tiers of cCRE-gene linkage, with level 6 the strongest evidence. A total of 401 genes from 29 GWAS loci were linked to lung cancer CCV-colocalizing cCREs at level 1 through 6 (**Table S10**). Among them were 64 linked genes from 18 loci with the strongest evidence at level 6 (**Table S11**). Nine of these 18 loci had one or more genes identified through bulk-tissue eQTL-based colocalization or transcriptome-wide association studies (TWAS) from published studies^9,43,44^, while 17 loci had one or more genes that have not been identified before (**Figure 4B; Table S12**). Notably, a CCV-colocalizing cCRE was linked to one to eight genes at level 6 (median 1 gene), and 82% of these cCREs were also linked to the same gene at level 4 or 5, indicating a redundant connection to target genes via modules of cCREs. Only 13% of the CCV-colocalizing cCREs were also located in the promoter of the same gene, suggesting that most of the CCV-colocalizing cCREs might regulate the target genes as distal enhancers. 74% of these putative distal enhancers were only detected in a single category of cell types. These data indicated that our cCRE-gene linkage approach could be validated by tissue-based eQTL methods and further identify additional candidate lung cancer susceptibility genes.

**Figure 4.**
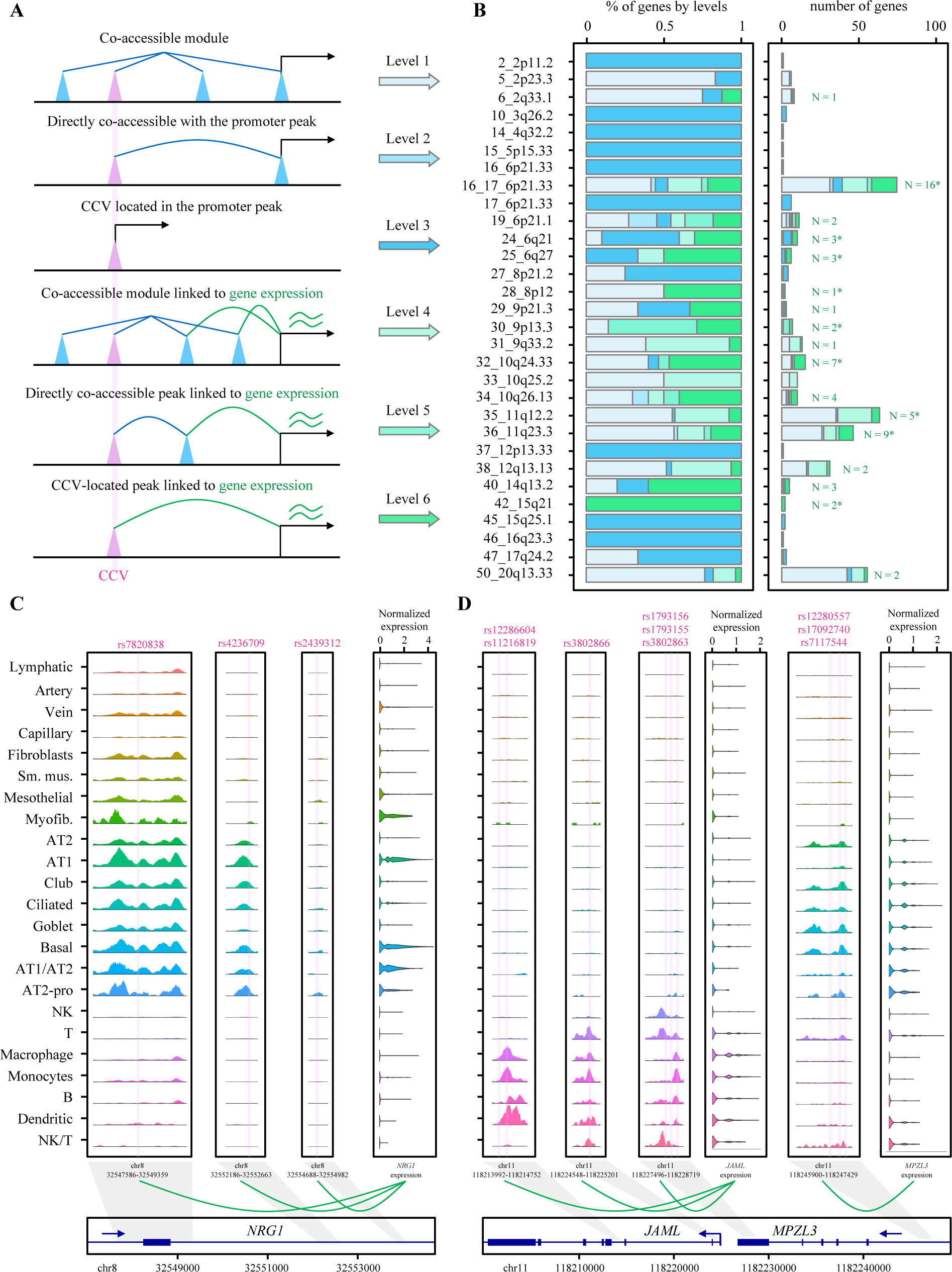
Linkage between cCREs and genes and representative loci showing context-specific genetic regulation mechanisms. **(A)** Illustration of the rationale in linking the cCREs to genes in different levels. cCRE module is represented by straight blue line. CCV-colocalizing cCREs are shown as pink cones. Highly co-accessible cCREs are represented by blue loop. The cCRE-gene correlations are represented by green loop. **(B)** The left panel presents the proportion of genes in different levels across GWAS loci. The right panel presents the number of genes in different levels across GWAS loci. The numbers refer to the level-6 gene numbers. *indicates genes identified by TWAS or colocalization from previous studies. **(C-D)** The rs IDs of the CCVs are presented in the upper part of the sequencing tracks of chromatin accessibility with their locations marked in the tracks using vertical lines. Each track represents the aggregate snATAC signal of all cell types normalized by the total number of reads in the TSS regions (normalized values for the left three ATAC tracks: 0-380; normalized values for the right four ATAC tracks: 0-240). Arrows show the transcriptional directions of the genes.

For example, an epithelial cell-specific CCV-colocalizing cCRE at 30_9p13.3 locus (**Figure S5**) was linked to *AQP3* based on level 4, 5, and 6 (**Table S10**). This locus was initially identified through a lung cancer TWAS^35,43^ using lung tissue eQTL data and later became genome-wide significant in a larger GWAS^36^. Over-expression of *AQP3* in cultured lung fibroblasts lead to increased endogenous DNA damage levels^43^. Our data validated *AQP3* as a lung cancer susceptibility gene in this locus and suggested potential involvement of epithelial-specific cCREs in its regulation. In the 6_2q33.1 locus, two CCV-colocalizing cCREs are located in *CASP8*. While a multi-cell-type cCRE (Chr2: 201,257,278-201,258,769) was linked to *CASP8* based on level 1, 2, or 3, an immune-cell-specific cCRE (**Figure S5**) was linked to *CLK1* expression by level 4, 5, or 6 (**Table S10**). *CLK1* is ∼400kb away from the linked cCRE and encodes Cdc2-like kinase 1 which is a splice factor kinase that has roles in tumorigenesis of multiple cancer types^45–47^. This locus is a multi-cancer-associated locus, and classically the role of *CASP8* in cancer cells has been mainly considered in their down regulation and consequent escape of apoptosis rather than immune-cell specific roles^48^. Our data suggested the presence of potential cell-type specific target genes of this locus.

### Coinherited CCVs are linked to multiple cell-type-specific target genes

In addition to identifying previously unknown candidate target genes from lung cancer GWAS loci, our dataset allowed us to deconvolute complex cell-type- and context-specific gene regulation patterns by multiple coinherited lung cancer-associated variants. First, we observed that multiple high-LD CCVs might individually affect the target gene expression via separate but correlated cCREs. In the locus 28_8p12, three CCVs colocalized with three different cCREs that are mainly observed in epithelial category. All three cCREs belong to the same cCRE module, and each of them individually was linked to *NRG1* expression at level 6 (**Figure 4C**). Consistent with this finding, these three cCREs overlapped with the region that was assigned to *NRG*1 based on the activity-by-contact chromatin interaction model^13^ in lung cell lines. *NRG1* was also a TWAS gene in this locus using GTEx lung tissues eQTL dataset. Notably, all three CCVs are in high-LD with the GWAS lead variant (*R*^2^ = 0.99 – 1, EUR), suggesting that multiple functional variants in LD might share the same target gene.

Second, we observed that distinct subsets of CCV-colocalizing cCREs could be linked to cell-type specific target genes. For example, the 36_11q23.3 locus includes two clusters of CCVs, among others, that are linked to two neighboring level-6 genes in a cell-type specific manner (**Figure 4D**). Namely, three cCREs colocalizing with six CCVs were mainly detected in immune-category cells and linked to *JAML* expression. Another cCRE colocalizing with three CCVs were mainly detected in epithelial-category cells and significantly correlated with *MPZL3* expression. Both of these cCREs groups include one or more CCVs with the evidence of footprints for allelic-binding TFs. Notably, both *JAML* and *MPZL3* have been identified by eQTL colocalization and TWAS approaches using lung tissues ^9,43^, but it has not been clear which genes are expressed and thus functional in what lung cell types or contexts. These data suggested that both epithelial and immune-cell specific lung cancer susceptibility genes could be functioning together based on the cellular contexts within the same lung cancer-associated locus.

## Discussion

In this study, we established a customized single-cell multiome dataset representing a lung cancer-relevant environmental exposure and presumed cell types of origin. Using this dataset, we identified candidate functional variants and susceptibility genes for most published lung cancer-associated loci, many of which displayed cell-type and context specificity. We also demonstrated the feasibility of using cryopreserved dissociated cells from solid tissues for single-cell applications by employing FACS-sorting and cell type balancing. Fresh tissue processing of surgically resected solid tissues is still a logistical challenge for many single-cell applications. While new technologies are emerging to use fresh-frozen or formalin-fixed paraffin-embedded tissues, our approach is an alternative that provides flexibility of cryopreserving dissociated tissues and enriching cell groups of interest by cell surface marker-based sorting.

Using single-cell multiome approach, we found that cCREs of lung tissue are mostly cell-type specific, and epithelial and immune cell groups contribute the most to genetic underpinning of lung cancer risk. Single-cell based TRS assessment highlighted AT2 cells and AT2-proliferating cells, among epithelial cell group, as the top contributing cell types to lung cancer susceptibility. This finding is consistent with the previous understanding of AT2 cells being an origin of lung adenocarcinoma and also aligns with the hypothesis that rare stem/progenitor populations responsible for alveolar regeneration might be important in lung tumorigenesis. In addition, we provide evidence for the long-regarded importance of immune cell populations in lung tumorigenesis. While a recent LD score regression analysis based on lung single cell expression data suggested the importance of AT2 cells in lung adenocarcinoma heritability^18^, previous LD score regression results using bulk tissues and cell-line based annotation datasets hinted at the roles of immune cells in lung cancer susceptibility, especially in European smokers^5,37^. Immune cell populations could interact with tumor cells originating from epithelial cells to promote their growth and selection through inflammatory signaling, for example, but they could also suppress tumor cell growth via immune surveillance during the trajectory of tumorigenesis. The potential interplay of epithelial and immune cell populations could be observed in multiple lung cancer loci; CCVs from 14 loci overlapped with both epithelial-specific and immune-specific cCREs. Multiple examples from cCRE-gene linkage also suggested that different subsets of CCVs might regulate epithelial or immune-cell specific target genes in a single locus.

CCV-cCRE colocalization enabled annotation of lung cancer GWAS loci based on rare or hard-to-culture cell types from normal lung tissues. For example, AT2 and proliferating sub-populations of AT2 cells are difficult to culture, making it challenging to investigate variant functionality for the genes that are specifically expressed in these cell types. Furthermore, we observed that long suspected roles of candidate genes aligned with their cellular contexts based on the cCREs specific to rare lung cell types. The 5p15.33 locus near *TERT* is one of the strongest lung cancer signals and associated with multiple cancers. While previous studies in the context of multiple cancer types identified functional variants regulating *TERT* expression, due to its inactivation in normal differentiated tissues, eQTL of *TERT* has not been detected in the GTEx lung tissues. By inspecting a rare stem-like cell population of lung epithelium, basal cells (1.8%), we show that a subset of CCVs in this locus overlaps with a cCRE distinct to this cell type. Due to the small number of cells in this population of cells, cCRE-gene linkage was not observed for *TERT* expression. However, one of the colocalizing variants displayed significant allelic transcriptional activity in A549 lung cancer cells with the risk-associated variant showing higher expression, suggesting that these variants might contribute to *TERT* expression.

Our dataset provided additional insights to potential mechanisms of multiple high-LD CCVs regulating their target genes in a context-dependent manner. While conventional approaches using bulk tissue or cell lines such as eQTL colocalization/TWAS and chromatin interaction analyses typically identify multiple target genes from a single GWAS locus, it is difficult to prioritize and interpret what genes are functional in what contexts. In our prime examples, we observed that some of these cases could be explained by cell-type specific gene regulation via different subsets of trait-associated variants. In the 36_11q23.3 locus, *JAML* expression was correlated with three different CCV-colocalizing cCREs in immune cell types while the cCRE are almost undetectable in epithelial cell types. Conversely, epithelial cell-specific CCV-colocalizing cCRE was linked to epithelial cell-preferential expressed genes including *MPZL3*, which is adjacent to *JAML*. *JAML* encodes a junctional adhesion molecule playing a role in defense against infection, tissue homeostasis and repair, and inflammation in the tissue-resident gamma-delta T cells at the epithelial barrier^49,50^, suggesting its potential roles in lung tumorigenesis through interplay with epithelial cells. We also show in the example of *NRG1* that multiple high-LD variants that might be indistinguishable in statistical approaches can be strongly associated with the same target gene while located in three separate cCREs. *NRG1* encodes Neuregulin-1, a ligand containing epidermal growth factor (EGF)-like receptor tyrosine kinase (RTK)-binding domain, and somatic fusion of *NRG1* is a rare tumor driver event but most frequently observed in lung adenocarcinoma cases among all cancer types^51^.

Our cell-cell communication analysis revealed an interesting phenomenon of MHC-I and MHC-II pathways displaying an opposite direction of changes between ever- and never-smokers. Anecdotal findings based on average gene expression levels of certain lung adenocarcinoma cell clusters were observed to similar effect in a previous study^33^, and MHC-II genes suppression was also shown in airway epithelial cells of smokers without reported lung cancer^29^. While the MHC-II communication reduction in ever-smokers seems prominent in myofibroblasts and ciliated cells, MHC-I communication reduction in never-smokers are similar across the cell types but slightly larger in AT2-proliferating and goblet cells. Together with the report that lower expression of MHC-I genes are observed in never-smoker lung adenocarcinoma cells^33^, our finding suggested that MHC-I communication reduction in never-smokers is already evident in normal lung cells including AT2-proliferating cells, and could lead to compromised antigen presentation and tumor cell surveillance by immune cells throughout the tumorigenesis. Multiple lung cancer GWAS hits were identified in the MHC loci on chromosome 6, and distinct signals were observed based on different studies representing European populations of mainly smokers or East Asian populations including a substantial proportion of never-smokers^53^. Our data suggested that smoking status-specific immune response through MHC pathways might contribute to lung cancer risk.

Our study has several limitations. First, the differentially expressed genes based on smoking status were mostly suggestive and did not pass the stringent cutoff. A larger study with well-documented exposure details could help improve the detection power. Second, our dataset does not represent all the detectable lung cell types that have been reported. It is possible that we lost specific epithelial cell types that are the most vulnerable (e.g., neuroendocrine cells). We also did not explore sub-populations of immune cells using sub-clustering. While we view that this is outside the scope of the current study, information on sub-populations of immune cell types could help interpret the immune-epithelial interplay underlying many lung cancer GWAS loci. Third, although very powerful in target gene identification, cCRE-gene linkage might include indirect association, especially for the connections with marginal p-values/scores and in longer-range. While indirect correlation could still provide valuable information, direct linkage could be established using further experimental validations.

In summary, our study provided a unique dataset to characterize cell-type and context specific genetic regulation underlying lung cancer susceptibility.

## Methods

### Human subjects and tissue collection

We collected tumor-distant normal lung parenchyma tissue samples from patients who underwent lobectomy with a curative aim for primary lung adenocarcinoma. Most of patients presented stage IA adenocarcinoma, and tumor stages were comparable between the groups based on sex or smoking status; in each of the four groups, >=75% of the patients presented stage I. Tumor-distant normal samples were obtained from the periphery of the lobe, more than 2 cm away from the tumor edge, which is unlikely to present tumor phenotype based on guidelines of the National Comprehensive Cancer Network. Dissociated tissues from four of the female never-smokers were combined to produce a single sample with a similar total cell counts to other samples, which resulted in a total of 16 samples (8 each from ever- and never-smokers) (**Table S1**). All the participating patients did not receive any other cancer treatment prior to surgery. The sample size of comparing 8 vs. 8 for assessing smoking effect was determined based on a recent scRNA-seq study that has successfully detected differential gene expression between smokers (n = 6) and non-smokers (n = 6) in human tracheal epithelium^29^. Never-smokers were defined as individuals who have smoked < 100 cigarettes in the lifetime. Ever-smokers were defined as individuals who have smoked ≥ 100 cigarettes in the lifetime, and their smoking status was recorded (age to start/quit smoking, current/former smoker, and self-reported pack years). There were no significant differences of age between ever- and never-smokers. Detailed characteristics of the patients included in the study are in **Table S1**. Patient tissues were obtained under a protocol approved by Yonsei University Health System, Severance Hospital, Institutional Review Board (IRB 4-2019-0447, 4-2022-0706), and informed consent was obtained from each patient prior to surgery. All experiments were performed following applicable regulations and guidelines.

### Tissue dissociation and cryopreservation

Fresh tissue dissociation and cryopreservation conditions were optimized to maximize cell viability and facilitate proper preservation of diverse cell types. Specifically, digestion enzyme titration was performed and freezing media were compared based on epithelial cell retention measured by flow cytometry after dissociation and freezing/thawing, respectively. The tissues were collected at a ∼2 cm^3^ size and put in MACS Tissue Storage Solution (Miltenyl biotec cat.130-100-008) within 30 mins of surgical dissection. Samples were kept at 4°C and processed within 3 hours of dissection. Tissues were chopped into 2 mm diameter pieces within a dissociation tube to reduce cell loss. Multi Tissue Dissociation Kit 1 and gentleMACS Octo Dissociator (Miltenyi Biotec) were used for dissociation of the tissue into single-cell suspension with a reduced amount of enzyme R (25% of the regular amount). Red blood cells were removed using Red Blood Cell Lysis Solution (Miltenyl biotec cat.130-094-183). Single-cell suspension of samples was frozen using 90% FBS and 10% DMSO and stored in liquid nitrogen until further processing except for the time of transfer in dry ice.

### Single cell processing and cell-type balancing

Given that immune cells were reported to be overrepresented in lung tissues^15^ and epithelial cells are relatively vulnerable to freezing and thawing, we employed balancing of major cell groups of the lung using cell surface marker labeling followed by FACS sorting. Balancing of fresh-dissociated lung cells using FACS sorting has been reported by other groups^15^. Specifically, the dissociated single-cell suspensions were thawed and filtered using a 70 μm cell strainer (Miltenyi Biotec) to remove debris. The filtered cells were labelled with cell viability marker (DAPI) and antibody markers of EPCAM, CD31, and CD45. Live single-cells (DAPI-negative) were sorted based on three gates: EPCAM^+^CD45^-^ (designated “epithelial”), EPCAM^-^CD45^+^ (designated “immune”), and EPCAM^-^CD45^-^ (designed “endothelial or stromal”). To enrich epithelial cells, which are considered to have key roles in lung cancer etiology, we collected all “epithelial” cells from EPCAM^+^CD45^-^ gates and balanced the ratios to 6:3:1 (“epithelial”: “immune”: “endothelial or stromal”).

### Nuclei isolation and single-nuclei multiome sequencing

Nuclei isolation was performed based on “Low Cell Input Nuclei Isolation” protocol (CG000365-Rev C, 10X Genomics) with a modification of the 1X lysis buffer treatment time (3-second duration). The concentration and treatment time of the lysis buffer were optimized based on the quality of nuclei measured by a 4-level grading of nuclei from microscopic images (non-lysed, ruptured, fair, intact) to achieve the minimum numbers of unlysed cells or ruptured nuclei and maximum numbers of intact and fair-quality nuclei. After processing all the samples following the optimized protocol, nuclei quality of all the samples was measured (∼200 nuclei/sample), and no significant differences were observed between samples from ever-vs. never-smokers (median percentage of fair or intact nuclei 57.5% vs. 56.5%). Isolated nuclei were used for single-cell capture and sequencing library preparation using the Chromium Next GEM Single Cell Multiome ATAC + Gene Expression Reagent Kits following the manufacturer’s guidelines (CG000338-Rev E, 10X Genomics, USA). The snATAC-seq and snRNA-seq libraries were sequenced on a NovaSeq6000 platform using the following cycles: snATAC-seq (Read 1N, 50 cycles; i7 Index, 8 cycles; i5 Index, 24 cycles; Read 2N, 49 cycles) and snRNA-seq (Read 1, 28 cycles; i7 Index, 10 cycles; i5 Index, 10 cycles; Read 2, 90 cycles). Each sample achieved > 499 million paired-reads for snATAC-seq and > 372 million paired-reads for snRNA-seq.

### Generating single-nucleus gene expression and chromatin accessibility matrices

Raw sequencing data were converted to fastq format using Cell Ranger ARC ‘mkfastq’ function (10x Genomics, v.2.0.1). snRNA-seq and snATAC-seq reads were aligned to the GRCh38 (hg38) reference genome using STAR and quantified using Cell Ranger ARC ‘count’ function (10x Genomics, v.2.0.1). The raw outputs from Cell Ranger ARC achieved 7,878 to 14,005 snATAC median high-quality fragments per nucleus, 98.1% to 98.9% snATAC valid barcodes, 1,905 to 2,941 snRNA median genes per nucleus, and 92.2% to 95.1% snRNA valid barcodes across 16 samples (**Table S2**). Gene expression and chromatin accessibility matrices were generated separately for each sample.

### Quality control and filtering

We used DropletQC (v.0.9) to remove “empty” droplets containing ambient RNA from the gene expression matrices^54^. Cell Ranger ARC-generated ‘possorted_genome_bam.bam’ files served as input for DropletQC. The parameter ‘nuclear_fraction’ was calculated using the function ‘nuclear_fraction_tags’ with default parameters. Empty droplets were identified by visualizing the density of nuclear fraction and settling the cutoff according to the “peak” in low-nuclear-fraction droplets (**Figure S6**). The resulting expression matrices were processed individually in R (v.4.1.3) using Seurat (v.4.0.6)^55^ and Gencode v.27 for gene identification. From each sample, we excluded cells with less than 500 or more than 25000 nCount_RNA (number of RNA read counts), cells with less than 1000 and more than 70,000 nCount_ATAC (number of ATAC read counts), and cells with more than 10% of counts corresponding to mitochondrial genes. In addition, Scrublet was applied to identify and remove doublets with an expected doublet rate 10% based on the loading rate^56^. Jointly applying ATAC and RNA filters resulted in a total of 117,911 cells from 16 samples with high-quality measurements across both modalities. After quality control and filtering, all cells across samples were merged.

### Gene expression data processing

Filtered gene–barcode matrices were normalized with the ‘SCTransform’ function of Seurat, and the top 2,000 variable genes were identified. Gene expression matrices were scaled and centered using the ‘ScaleData’ function.

### Peak calling and annotation

The snATAC peak calling and annotation were performed following the Signac pipeline^57^. Specifically, peaks were called using MACS2 and default parameters after combining the reads of all the cells in each cell type to determine the genomic regions enriched for Tn5 accessibility from snATAC fragments. Peaks were then annotated according to distance to protein-coding genes using ChIPseeker, with “tssRegion” setting of -3000 to 3000^58^. Term frequency inverse document frequency (TF-IDF) normalization on a matrix was performed using ‘RunTFIDF’ function.

### Clustering and cell-type annotation

Using the normalized gene expression data, we performed principal component analysis (PCA) with 50 PCs to compute and store. A uniform manifold approximation and projection (UMAP)-based approach was applied for expression matrices with the first 50 PCs and for chromatin accessibility matrices with the 2^nd^ through 50^th^ PCs (the first PC was excluded as this is typically correlated with sequencing depth). Both expression and chromatin accessibility matrices were corrected for batch effect using Harmony^59^. A Weighted Nearest Neighbor (WNN) method^55^ was applied to integrate the weighted combination of RNA and ATAC-seq modalities. The ‘FindClusters’ function was applied for clustering using smart local moving (SLM) algorithm for modularity optimization at a resolution of 0.5. Clusters were annotated based on canonical marker genes reported by three independent studies^15,18,22^. Clusters expressing canonical marker genes from two or more different cell types were designated as putative doublets and excluded, after which re-clustering was performed using the same parameters. Clusters with no detected marker genes were also excluded, after which the dataset was also re-clustered. Clusters in the final dataset representing subpopulations of the same cell type were grouped together for downstream analyses.

### Analysis of smoking-responsive genes

Smoking-responsive genes for each cell type were inferred by pseudobulk differential gene expression analysis using DESeq2 (v1.41.1). We initially performed a pilot analysis using a method without adjusting within-sample correlation (e.g., Wilcoxon rank-sum test) and observed p-value inflation in highly-expressed genes which was confirmed by a label-swapping and permutation analysis. To avoid false positive findings in highly expressed genes and also be able to incorporate other covariates in the model^60^ we chose a pseudobulk-based method. DESeq2 model assumes a negative binomial distribution^61^, and the gene expression counts were aggregated for each sample before differential expression analysis to account for the within-sample cell-cell correlation. Gender was incorporated as the covariate into the model. Given that the sample size is limited (8 ever-smokers versus 8 never-smokers) and the method is highly stringent, we report suggestive smoking-responsive genes using a relaxed cutoff of *P* < 0.0001 before multiple-testing correction.

### Intercellular communication analysis

CellChat (version 1.6.0) was used to infer ligand–receptor interactions based on scRNA-seq data^62^. CellChat applies a signaling molecule interaction database (CellChatDB.human) to predict intercellular communication patterns based on differentially overexpressed receptors and ligands, including soluble agonists/antagonists as well as membrane-bound receptors/co-receptors. Known ligand–receptor pairs from the CellChatDB database were used to compute the communication probability at the pair level and then summarized into interaction strength at the pathway level and across all the pathways. The intercellular communication analyses were performed separately using cells from ever-smokers and never-smokers for comparison at different levels. We nominated the top 5 pathways showing the largest difference in the relative sum of communication strength of all cell types (scale of 0 to 1) between ever- and never-smoker groups in each direction (top 5 increased and top 5 decreased in ever-smokers).

### Lung cancer GWAS candidate causal variants (CCVs)

A total of 51 genome-wide significant loci (**Table S7**) were included from 4 recent lung cancer GWAS studies: Mckay_2017 (European population^35^), Dai_2019 (European and East Asian population^36^), Byun_2022 (meta-analysis of European, East Asian, and African population^9^), and Shi_2023 (East Asian population^37^). A total of 2,574 CCVs were selected based on the following criteria (**Table S8**):

- For Mckay_2017, variants with log likelihood ratio (LLR) < 1:1000 with the primary lead SNPs based on the GWAS *P* values or *R*^2^ > 0.8 with the primary lead SNPs (1000 Genomes, phase 3, EUR population).
- For Dai_2019, variants with *R*^2^ > 0.8 with the primary lead SNPs (1000 Genomes, phase 3, EUR or EAS population) in 5 new loci.
- For Byun_2022, variants with LLR < 1:1000 with the primary lead SNPs based on the GWAS *P* values or *R*^2^ > 0.8 with the primary lead SNPs (1000 Genomes, phase 3, EUR, EAS, AFR, or ALL population) for 6 novel loci.
- For Shi_2023, variants with LLR < 1:1000 with the primary lead SNPs based on the GWAS *P* values or *R*^2^ > 0.8 with the primary lead SNPs (1000 Genomes, phase 3, EAS population) for 13 significant loci from the discovery stage. Variants with *R*^2^ > 0.8 with the primary lead SNPs for 9 loci significant in replication stage were included. Three independent SNPs from conditional signals were included, and they did not have proxy variants of *R*^2^ > 0.8 in EAS.

### Trait relevance score calculation

We inferred the lung-cancer-associated score for each cell using SCAVENGE pipeline^38^, based on snATAC-seq and lung cancer GWAS data. For GWAS input, we used flat probability scores evenly divided across all the CCVs in each locus. The peak-by-cell matrix of snATAC-seq data were processed using ArchR package^63^. gchromVAR was performed to calculate the original colocalization scores between GWAS and snATAC-seq. Seed cells proportion was set as 5%, and network propagation was applied to calculate the lung-cancer-associated score for each cell.

### Cell-type and category specificity of the accessible peaks

The specificity of the accessible peaks were assigned if they were overlapped with a single cell type or cell types in a single category (epithelial, immune, endothelial, or stromal). If a peak overlaps with cell types from multiple categories, we tested whether this peak can be mainly attributed to a single category by comparing the observed fraction of cell types to the expected 25^th^ and 75^th^ percentile in each category. For instance, if a peak overlaps with 7 epithelial, 2 immune, 1 stromal, and 1 immune cells, this peak is above the expected 75^th^ percentile of epithelial (> 6, considering 8 epithelial cells in total) but not in other three categories and thus is assigned to epithelial category. Peaks were assigned to multiple categories if they showed same levels of expected percentile in more than one category.

### Allelic effects of predicted TF binding, cell-type-specific TF abundance assessment, and TF footprinting

Prediction of variant effects on TF binding sites was performed with the motifbreakR package^64^ and a comprehensive collection of human TF-binding site models (HOCOMOCO, v11)^65^. We selected the information content algorithm and used a threshold of 10^4^ as the maximum p value for a TF-binding site match in motifbreakR^64^. The allelic-binding effect was defined by the difference between alternative allele score and reference allele score larger than 0.7. “Abundant TF” in a given cell type was defined as: 1) TF is expressed in > 50% of the cells in that cell type and 2) TF expression level in the same cell type is above 75 percentile of the values from all predicted TF-cell type pairs. TF footprint analysis was performed for each allelic-binding TF using the ‘Footprint’ function in Signac by restricting to the peak regions. A significant footprint was determined by visualizing the height of the motif-flanking region accessibility compared to the background (expected Tn5 insertion rate)^57^.

### Reporter assays for variant allelic function test

Allelic transcriptional activity of CCVs from the 15_5p15.33 locus was assessed as part of MPRA. MPRA library construction, transfections, sequencing, and data analyses methods were based on a previous study^42^. Briefly, MPRA libraries were generated by cloning 145bp genomic sequence encompassing each test variant with reference or alternative allele in forward and reverse complementary direction in front of the minimal promoter of the luciferase constructs. Each test sequence was associated with 25 unique sequence tags (12bp) at the 3’ untranslated regions of the luciferase gene. MPRA libraries were transfected to two lung cancer cell lines, A549 (lung adenocarcinoma) and H520 (lung squamous cell carcinoma) with or without short-term exposure (10 µM for 18 hr and 42 hr for A549 and H520, respectively) to a tobacco carcinogen, BaP. A linear regression was performed to assess the effect of allele on the transcript output measured by normalized tag counts across 25 different tags and 5 transfection replicates in each condition using the formula: Ratio = Allele + Strand (forward or reverse) + Transfection batch. To determine the significance of allelic difference, robust sandwich type variance estimate in the Wald test was performed.

### cCRE-module, cCRE-cCRE, and cCRE-gene correlation

The co-accessible cCRE modules of two or more cCREs were identified by Cicero with Louvain community detection algorithm and co-accessible score cutoff of 0.32 (automatically defined)^66^. A more stringent co-accessible score cutoff of 0.5 was used to define “directly co-accessible” cCREs. For cCRE-gene correlation, we identified cCREs that may regulate a given gene by computing the correlation between gene expression and accessibility at nearby cCREs, and correcting for bias due to GC content, overall accessibility, and peak size. Specifically, we performed the cCRE-gene Pearson correlation analysis by running the ‘LinkPeaks’ function with ‘distance’ of 10^6^ (+/- 1Mb of TSS), ‘min.cells’ of 10, ‘pvalue_cutoff’ of 0.05, and ‘score_cutoff’ of 0.05. A six-level target gene assignment for the lung cancer GWAS loci was performed based on the following criteria:

- Level 1 (module correlation with a promoter): CCV-colocalizing cCRE is a member of a cCRE module, and one or more member(s) is an annotated promoter cCRE
- Level 2 (direct correlation with a promoter): the accessibility of CCV-colocalizing cCRE is directly correlated (co-accessible score > 0.5) with the accessibility of an annotated promoter cCRE
- Level 3 (a promoter cCRE): CCV-colocalizing cCRE is annotated as a promoter
- Level 4 (module correlation with a gene expression): CCV-colocalizing cCRE is a member of a cCRE module, and the accessibility of one or more member(s) of the same module is linked to gene expression levels
- Level 5 (direct correlation with a gene expression-linked cCRE): the accessibility of CCV-colocalizing cCRE is directly correlated with that of a gene expression-linked cCRE
- Level 6 (a gene expression-linked cCRE): the accessibility of CCV-colocalizing cCRE is linked to gene expression levels

## Data and Code Availability

All single-cell sequencing raw and processed data will be deposited in the Gene Expression Omnibus (GEO) database and publicly available upon publication. Any additional information required to reanalyze the data reported in this paper is available from the lead contact upon request.

## Acknowledgements

This work has been supported by the Intramural Research Program (IRP) of the Division of Cancer Epidemiology and Genetics, National Cancer Institute (NCI), US National Institutes of Health (NIH). This work utilized the Biowulf cluster computing system at the NIH. We thank the members at the NCI Cancer Genomics Research Laboratory (CGR) and Center for Cancer Research Sequencing Facility (CCR-SF) for help with sequencing efforts and members of NHLBI flow cytometry core for help with FACS sorting. This work was supported by the National Research Foundation of Korea (NRF) grant funded by the Korean government (Ministry of Science and ICT) (No. 2023R1A2C1004922) awarded to Eun Young Kim. Erping Long is supported by the National Natural Science Foundation of China (Excellent Youth Scholars Program and 82090011) and Chinese Academy of Medical Sciences Innovation Fund (2023-I2M-3-010). Jun Xia is supported by the State of Nebraska LB595, LB692, and NIH/NIEHS R00ES033259 awards. We thank Michelle Antony and Elelta Sisay for proofreading the manuscript and Hangnoh Lee for helpful advice on the analyses. The content of this publication does not necessarily reflect the views or policies of the US Department of Health and Human Services, nor does the mention of trade names, commercial products, or organizations imply endorsement by the US Government.

## Declaration of Interests

The authors declare no competing interests.

---

**Figure S1.**
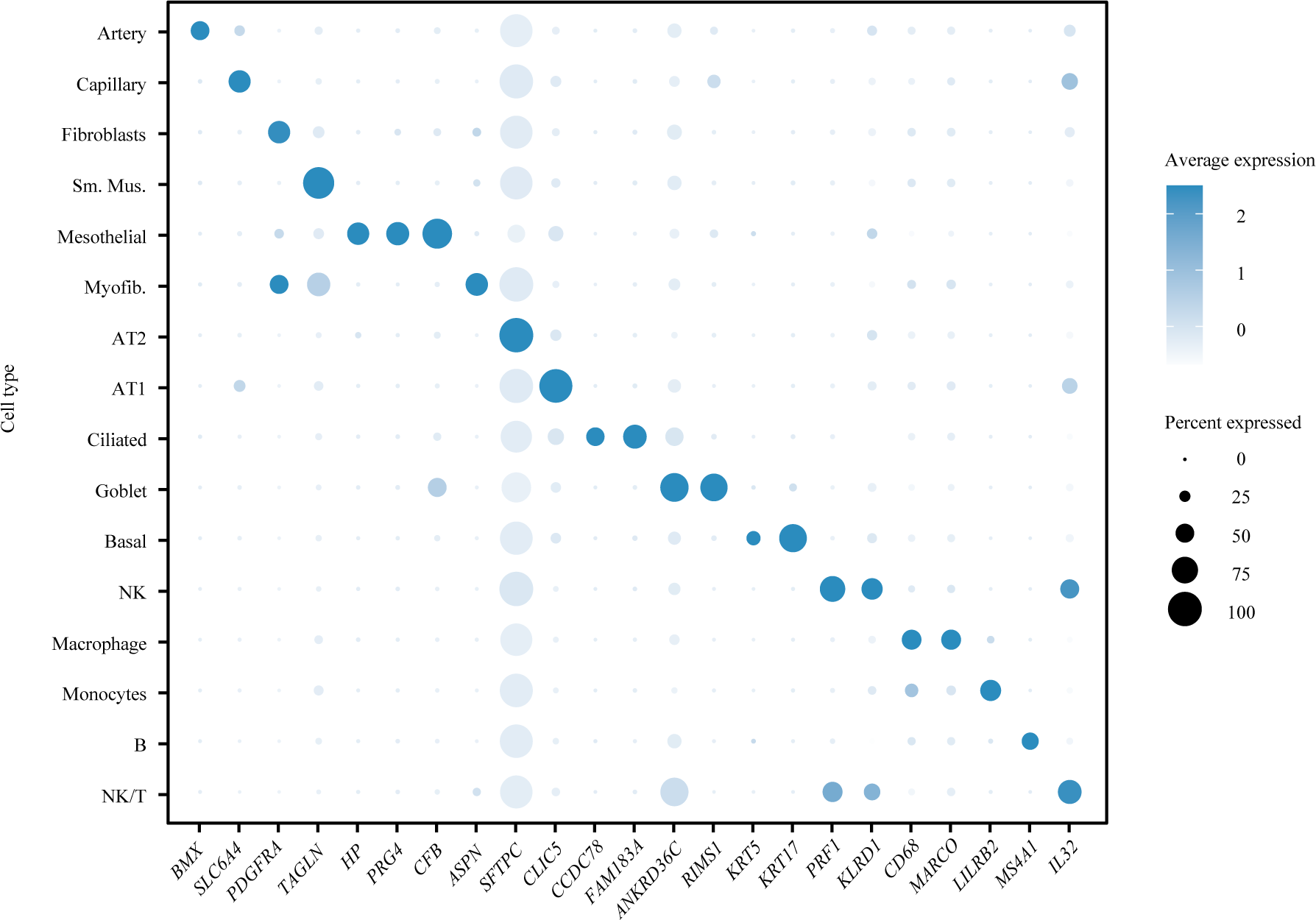
Additional canonical markers for cell-type annotation. Dot plot visualizing the normalized RNA expression of selected marker genes by cell type. The color and size of each dot correspond to the scaled average expression level and fraction of expressing cells, respectively.

**Figure S2.**
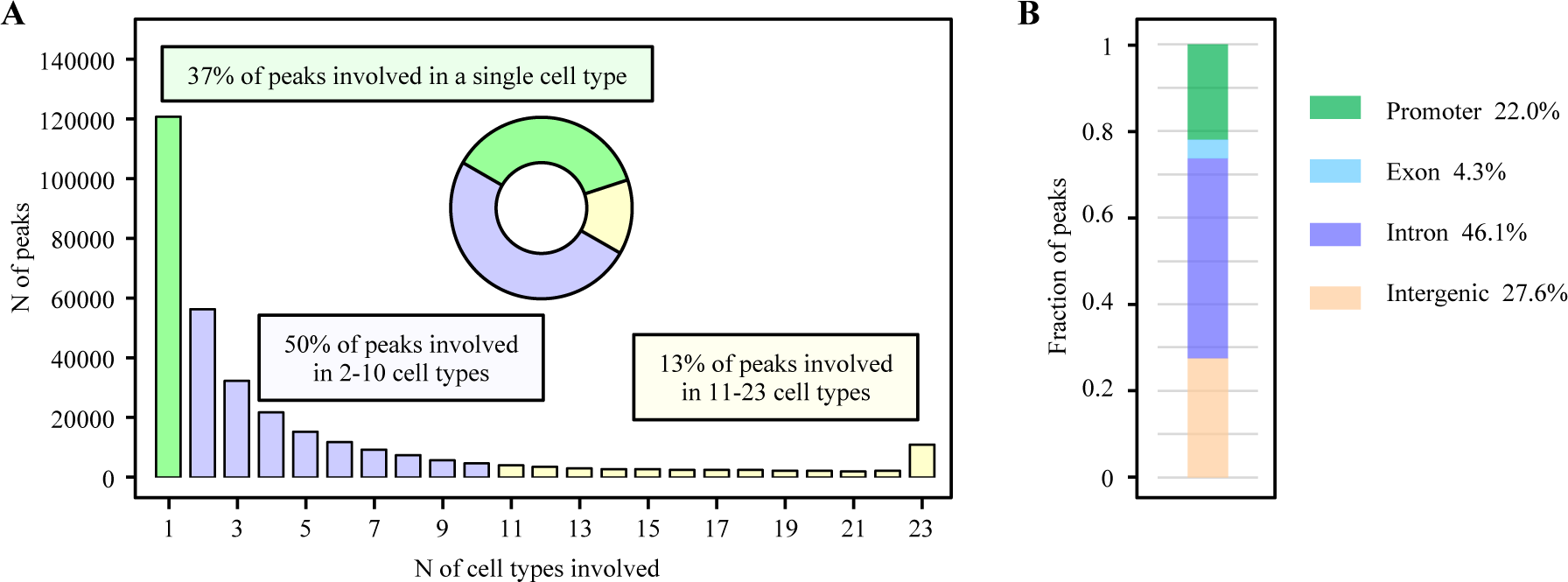
Accessible chromatin peaks identified from snATAC-seq data across different cell types. **(A)** Y-axis refers to number of peaks and X-axis refers to number of cell types involved. The piechart represents the proportion of peaks in single, 2-10, or 11-23 cell types. **(B)** Fraction of peaks were annotated into different categories (promoter, exon, intron, or intergenic).

**Figure S3.**
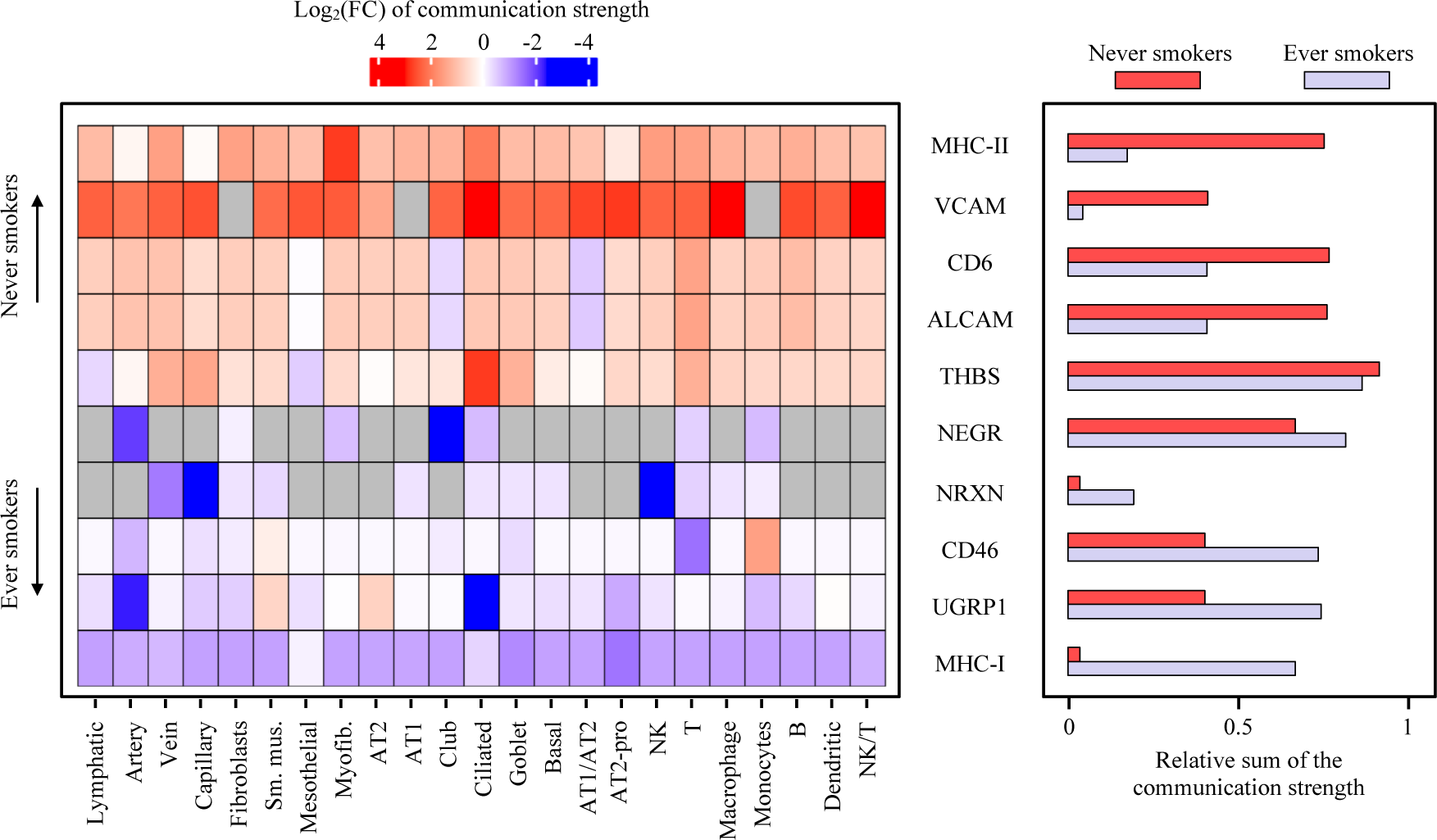
Differential intercellular communication based on smoking status. Heatmap of the top 5 elevated pathways in ever-(blue) and never-smokers (red) across cell types are shown on the left. Color indicates the log-transformed ratio of pathway-level communication strength between ever- and never-smokers (relative to never-smokers). Gray indicates that the pathway strength is 0 in the cell types of smokers and never smokers. The right part presents the summarized communication strength of each pathway for ever-(blue) and never-smokers (red).

**Figure S4.**
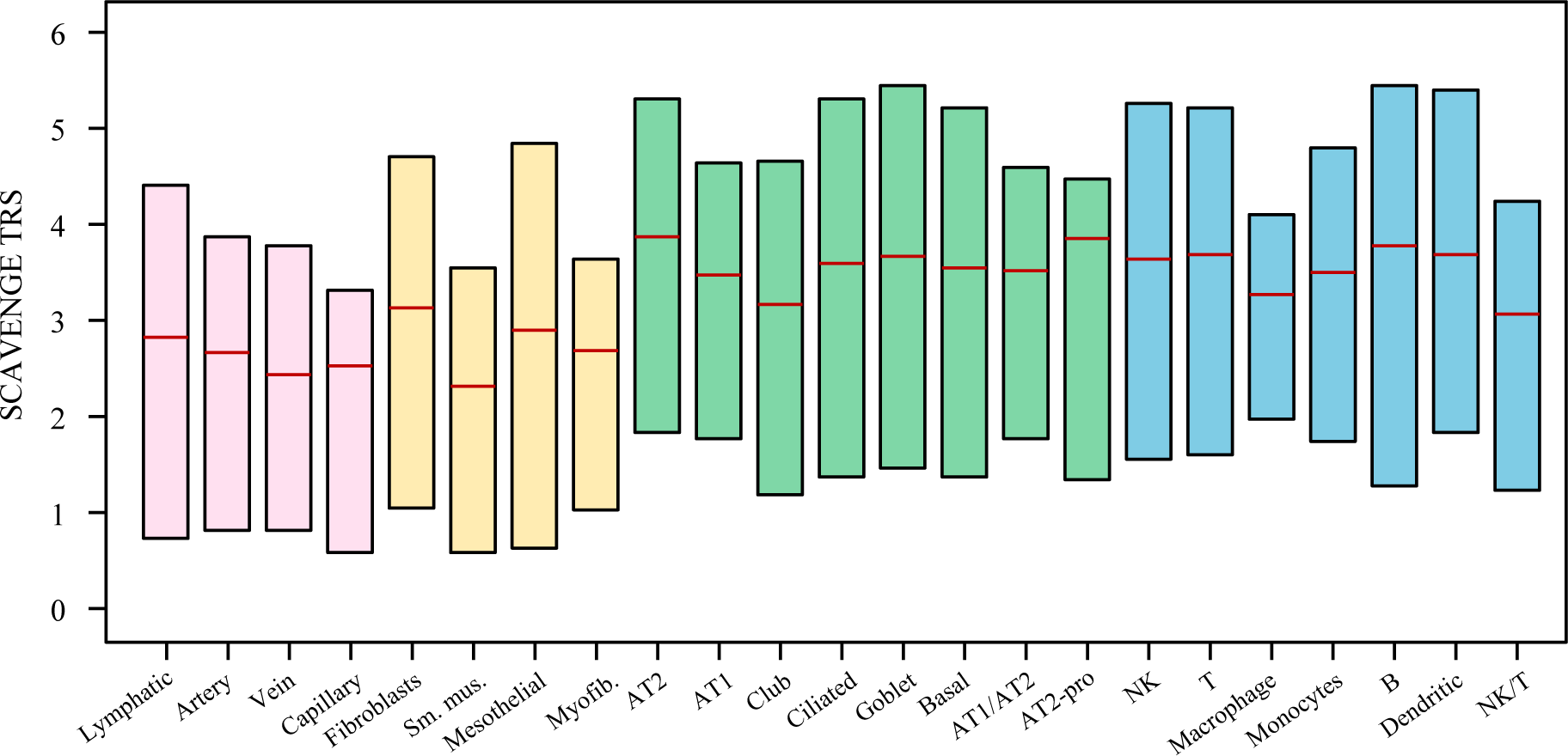
Trait relevance scores across cell types. The distribution of trait relevance scores (TRS) of lung cancer risk across cell types (endothelial in red, stromal in yellow, epithelial in green, and immune in blue) were presented as a boxplot. The red horizonal band shows the mean score, and the box indicates the middle 50% of cells.

**Figure S5.**
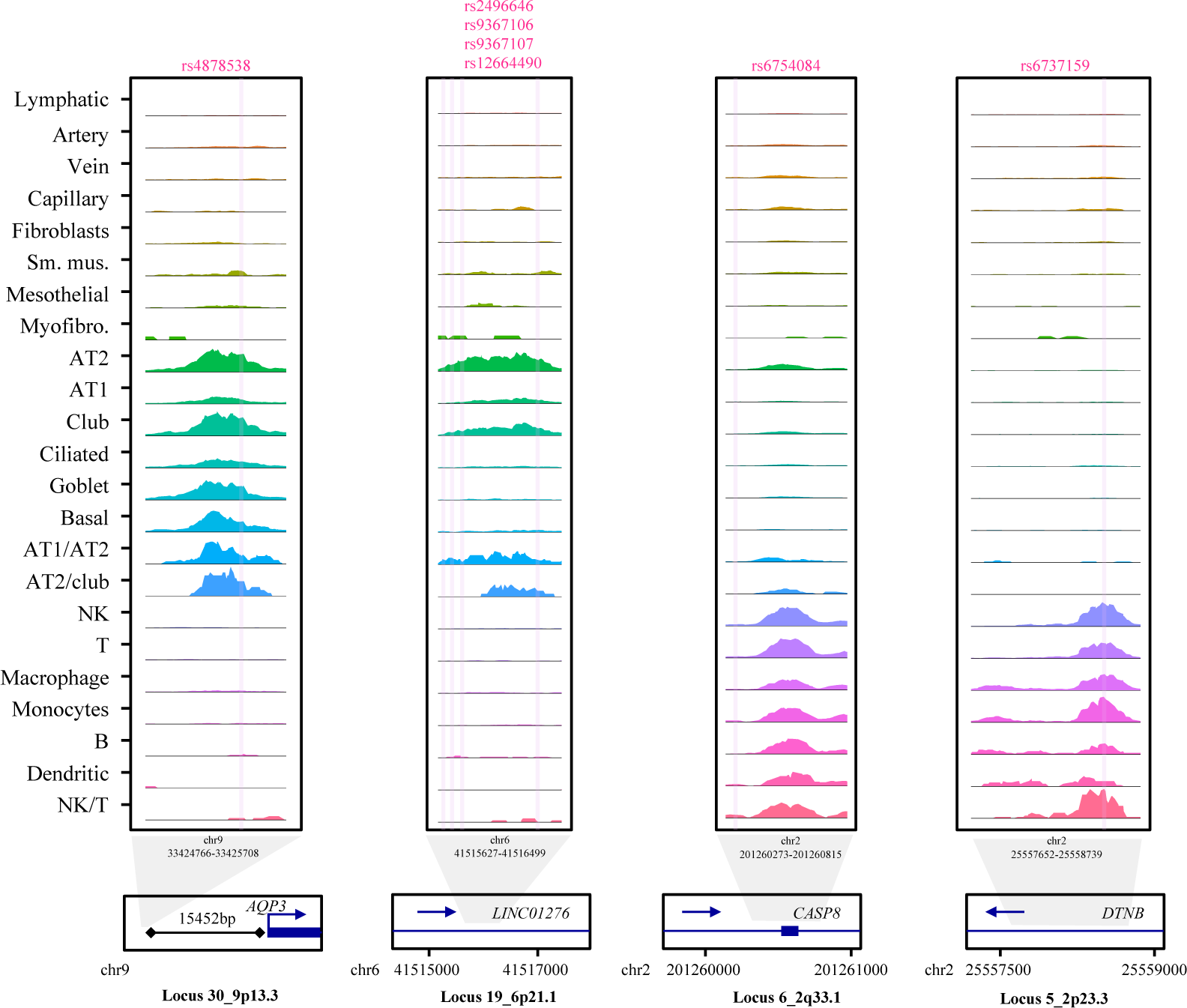
Epithelial or immune cell specific CCV-overlapping cCREs. The sequencing tracks representing chromatin accessibility of four different loci are displayed (locus IDs at the bottom). The rsIDs of CCVs are shown above the tracks and marked with vertical pink lines to indicate their genomic positions. Each track represents the aggregated snATAC signal of all cell types, normalized by the total number of reads in the regions (normalized values from left to right: 0-180, 0-180, 0-500, 0-240). Arrows depict the transcriptional directions of each gene.

**Figure S6.**
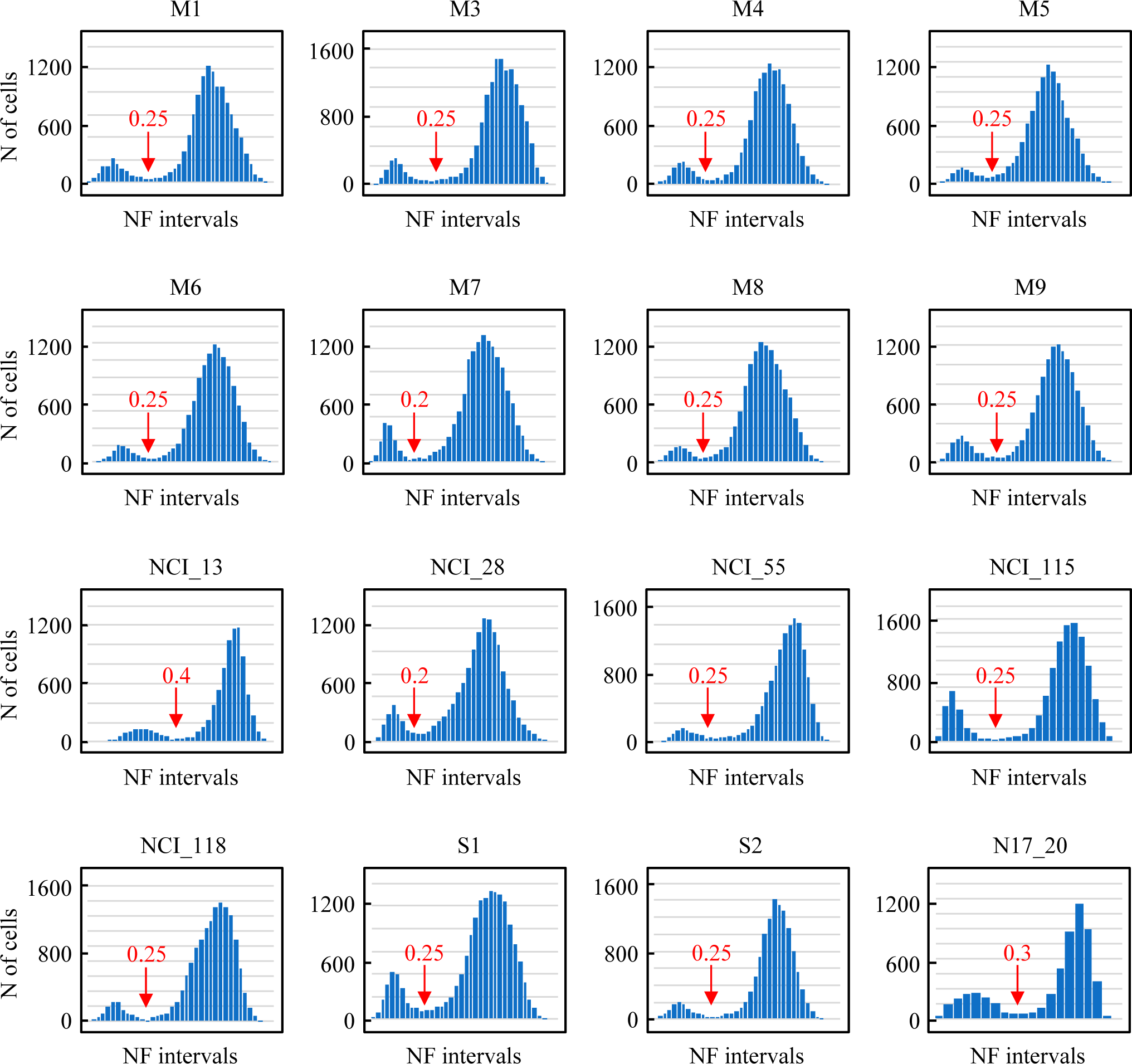
Determination of the cutoff in defining the empty droplet using DropletQC. The nuclear fraction (NF) of each cell is calculated by DropletQC. Empty droplets were identified by visualizing the density of nuclear fraction and setting the cutoff according to the “peak” in low-nuclear-fraction droplets.

## Notes

### Competing Interest Statement

The authors have declared no competing interest.

